# Vinorelbine enhances the efficacy of GM-CSF-armed oncolytic vaccinia virus in a preclinical model of ovarian high grade serous carcinoma

**DOI:** 10.1101/2025.02.04.636413

**Authors:** Stephanie Drymiotou, Christophe J. Queval, Katherine E. Tyson, Lesley A. Sheach, Antonio Postigo, Ilaria Dalla Rosa, Darren P. Ennis, Michael Howell, Iain A. McNeish, Michael Way

**Author notes:** Current address: Department of Obstetrics & Gynaecology, Barts Health NHS Trust, London, E11 1NR, UK. Current address: Stratosvir Limited, Stevenage, SG1 2FX, UK.

## Abstract

Vaccinia virus, known for its clinical safety has a tropism for primary and metastatic tumours as well as ovarian tissue. Consequently, oncolytic approaches with recombinant vaccinia viruses have emerged as attractive agents against ovarian cancer. Unfortunately, oncolytic vaccinia monotherapies are yet to live up to their potential promise. Given this, there is a need to identify combination agents that improve the effectiveness of vaccinia in ovarian cancer treatment. We screened 9,000 compounds to identify drugs that enhance the ability of a recombinant vaccinia virus lacking VGF and F1 (ΔVF) to induce death of ID8 *Trp53^-/-^* murine ovarian cancer cells. We identified a class of tubulin polymerisation inhibitors including vinorelbine. The combination of vinorelbine and vaccinia induces ID8 *Trp53^-/-^* cell death via apoptosis. In a syngeneic mouse model of high grade serous ovarian carcinoma, ΔVF virus lacking the viral thymidine kinase (TK), armed with GM-CSF and expressing NeonGreen (ΔVFTK-NG-GM-CSF) is tumour specific. A combination of the ΔVFTK-NG-GM-CSF virus with vinorelbine prolongs mouse survival compared to the treatment of mice with either agent alone. Our study suggests vinorelbine is a promising agent to combine with oncolytic vaccinia virus approaches for the management of ovarian cancer.

## Introduction

Ovarian cancer (OC) is the sixth most common female malignancy in the UK and the leading cause of death from a gynaecological cancer.^1^ It is a highly heterogeneous disease with histotypes varying in their genetic and molecular profiles, clinical phenotypes and risk factors.^2^ Despite its heterogeneity, standard management has remained the same for decades and includes surgical debulking combined with platinum-taxane chemotherapy.^3^ Response rates are initially high, especially in high grade serous carcinoma but 80% of patients with advanced disease recur, with all relapsed disease ultimately developing fatal therapy resistance.^4^ The introduction of bevacizumab (VEGF inhibitor) and PARP inhibitors in treatment regimens based on tumour molecular biomarker status can extend progression-free survival.^5,6^ However, the failure of maintenance therapies to extend overall survival for the majority of patients highlights the need for new therapies that overcome chemoresistance.^7,8^

Vaccinia virus represents a promising oncolytic agent for OC due to its natural tropism for ovarian tissue.^9^ The virus is best known for its use as the vaccine for smallpox eradication in the 1980s, demonstrating its safety and tolerability in humans.^10^ It also preferentially infects, replicates and kills primary and metastatic tumours over normal tissues.^11,12^ This tumour-specific cytotoxicity reduces the systemic side effects induced by the virus compared to existing chemotherapeutics.^13^ It also exclusively replicates in the cytoplasm thus avoiding DNA insertions into the host genome.^14^ Moreover, the safety and tumour specificity of the virus has been further enhanced through genome manipulation, by deleting genes encoding for virulence factors and proteins regulating nucleotide metabolism.^15^ For example, loss of thymidine kinase (TK) and vaccinia growth factor (VGF) significantly reduces viral pathogenicity, reducing side effects in patients, while simultaneously increasing tumour specificity.^16,17,18^ Therefore, TK and VGF are deleted from most vaccinia strains used in clinical trials.^19^ Vaccinia can tolerate 25-40 kb foreign DNA insertions and transgenes have been expressed in non-essential loci to improve its oncolytic properties and immunogenicity.^20,15^ Vaccinia kill cells by direct oncolysis and induces immunogenic cell death, an attractive feature that can be exploited and enhanced when used as an oncolytic virus against OC tumours.^21,22^

Olvi-Vec, derived from the Lister vaccinia strain, is the only oncolytic that has progressed to a phase III clinical trial for the treatment of platinum resistant OC in combination with chemotherapy and bevacizumab.^23,24^ This trial provided evidence that patients with platinum-resistant disease can respond to platinum chemotherapy. In contrast, a phase II clinical trial using the attenuated modified vaccinia Ankara virus expressing 5T4 tumour associated antigen (MVA-5T4, Trovax) monotherapy in asymptomatic women with recurrent ovarian cancer failed to show a significant improvement compared to placebo.^25^ This trial highlights the importance of combination therapies for OC with vaccinia.

The aim of our study was to identify a combination regimen that enhances the efficacy of vaccinia in treating ovarian carcinoma *in vivo.* We took advantage of a recombinant vaccinia virus lacking F1, a viral inhibitor of apoptosis ^26,27^ and the vaccinia growth factor (VGF).^28,29^ The recombinant ΔVF virus induces increased cell death during infection compared to the parental Western Reserve (WR).^30^ This is the first study in which the oncolytic efficacy of ΔVF was assessed *in vivo* in a syngeneic high grade serous ovarian carcinoma mouse model. Additional recombinant viruses have been constructed to improve the tumour specificity and immunogenicity of ΔVF. A high throughput compound screen was also conducted to identify combination partners that enhance the efficiency of ΔVF and its derivatives in killing ovarian cancer cells and inhibiting tumour progression.

## Results

### Targeting and arming ΔVF recombinant virus

To improve the potential tumour specificity of the ΔVF virus, we deleted the thymidine kinase (TK) gene to generate the ΔVFTK virus. To facilitate visualisation of infected cells, we also introduced NeonGreen (NG) into the TK locus, generating the ΔVFTK-NG virus. This virus was additionally armed with granulocyte-macrophage colony-stimulating factor (GM-CSF) to enhance its immunogenicity (ΔVFTK-NG-GM-CSF). Characterisation of the new recombinant viruses demonstrated they have similar viral protein expression and spread profiles to the ΔVF virus (Figure 1A and B). Immunoblot analysis confirmed NeonGreen (26 kDa) and the NeonGreen-GM-CSF fusion protein (47 kDa) were expressed in cells infected with the ΔVFTK-NG and ΔVFTK-NG-GM-CSF viruses (Figure 1A). Moreover, all viruses retained the pro-apoptotic activity of the parental ΔVF virus (Figure 1C).

**Figure 1.**
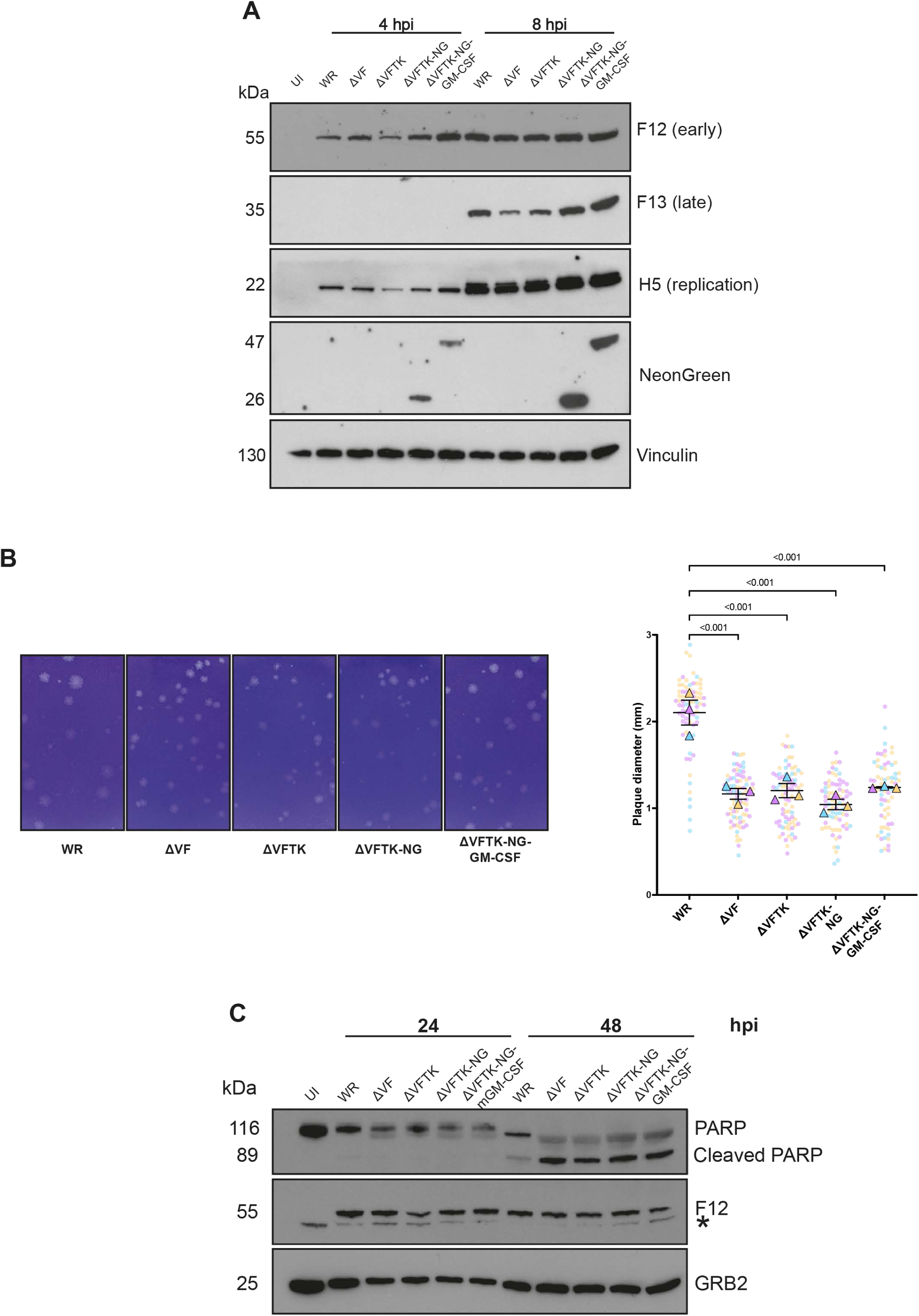
Characterisation of recombinant vaccinia viruses. **A.** Immunoblot analysis comparing the expression levels of the indicated viral proteins in HeLa cells infected with the specified viruses. Vinculin is the cell loading control. The experiment was repeated three times and a representative example is shown. **B.** Representative images of plaque formation by the indicated virus strains in BS-C-1 cells at 72 hpi. The graph represents quantitative analysis of plaque diameter measured in mm for the indicated viral strains. The three independent plaque assays are represented by the three different colours in the SuperPlot. Each dot represents one plaque, and the triangles represent the medians of all the plaques measured in each experiment. Data are represented as mean ± SD. One-way ANOVA was used to determine significance between all groups with Tukey multiple comparisons post-hoc test. **C.** Immunoblot analysis assessing PARP cleavage in ID8 *Trp53^-/-^* cells infected with the indicated viruses. F12 and GRB2 represent viral and cell loading controls respectively. The asterisk (*) indicates a non-specific band. Hpi = hours post infection. The experiment was repeated three times and a representative blot is shown.

The oncolytic efficacy of the new recombinant viruses was examined *in vivo* using a syngeneic mouse model of high grade serous ovarian carcinoma (ID8 *Trp53^-/-^*) (Figure 2A). The heat inactivated ΔVF (ΔVFi) virus was used as vehicle and the median survival of mice in this group was 45.5 days. The median survival of mice inoculated with ΔVF, ΔVFTK, ΔVFTK-NG and ΔVFTK-NG-GM-CSF viruses was 51.5, 51, 51 and 54 days, respectively. Log-rank analysis demonstrated that only ΔVFTK-NG-GM-CSF (p = 0.005) significantly prolonged mouse survival (Figure 2B) compared to vehicle control. Viral inoculation of mice with the recombinant viruses had no effect on the size of their liver and spleen nor ascitic volumes (Figure 2C). NeonGreen expression did not affect mouse survival as mice inoculated with ΔVFTK and ΔVFTK-NG had the same median survival of 51 days. An additional survival experiment in which viral treatment began 14 days post intraperitoneal (IP) ID8 *Trp53^-/-^* cell injection with the mice receiving four viral IP doses instead of three was conducted. In this protocol, both ΔVFTK-NG and ΔVFTK-NG-GM-CSF viruses significantly prolonged mouse survival compared to the heat inactivated ΔVF control (Figure S1). These data, provide further support for the contribution of viral oncolysis to the observed therapeutic benefit and indicate that the ΔVFTK-NG-GM-CSF was the only recombinant virus that consistently improved survival. These experiments also illustrated the need to identify combination agents to improve the efficacy of vaccinia virus in the treatment of ovarian cancer.

**Figure 2.**
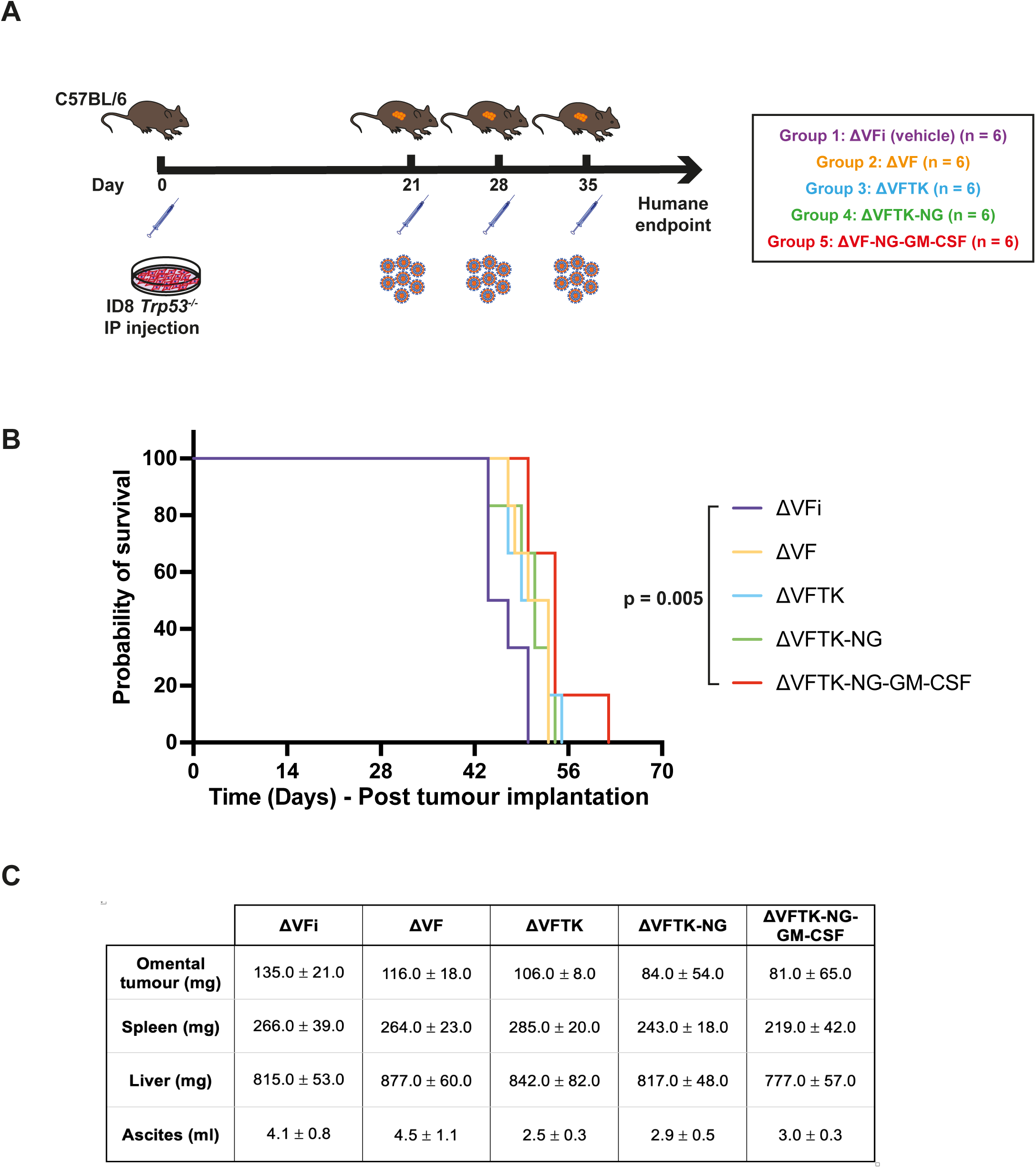
Efficacy of recombinant vaccinia viruses in vivo. **A.** Schematic representation of the experimental design of the *in vivo* survival study. Five groups of mice were injected IP with ID8 *Trp53^-/-^* cells on day 0 and subsequently inoculated with the indicated viruses on day 21, 28 and 35. ΔVFi is the control heat inactivated virus. **B.** Kaplan-Meier survival curve showing survival data for each virus analysed by log-rank test. **C.** Quantification of omental tumour, spleen and liver weights as well as ascitic volumes for each group. Data are represented as mean ± SD. One-way ANOVA was used to determine significance between all groups with Tukey multiple comparisons post-hoc test.

## Identifying a combination agent to improve the efficacy of oncolytic vaccinia

A high throughput drug screen was conducted on ΔVFTK-NG infected murine ovarian cancer (ID8 *Trp53^-/-^*) cells (Figure 3A). The library consisted of 9,000 well-characterised compounds that are in clinical development or have been approved by Food and Drug Administration (FDA) and/or European Medicines Agency (EMA) and used in clinic. NeonGreen fluorescence was used to identify infected cells and DAPI staining of nuclei was used to calculate the number of cells remaining at the end of the screen. After imaging of wells compounds were allocated into one of four categories (‘no effect’, ‘kill infected cells’, ‘block virus replication’ and ‘toxic’) based on the number of infected (green) and uninfected (non-green) cells relative to infected DMSO controls on the same plate (Figure 3B). The ‘no effect’ are wells with similar numbers of infected and uninfected cells compared to the DMSO control, whereas ‘block virus replication’ are those with reduced infected and increased uninfected cells due to cell proliferation. Toxic compounds resulted in a significant loss of infected and uninfected cells. The ideal compounds are the ‘kill infected cells’ category as they represent wells with fewer infected but similar numbers of uninfected cells compared to the DMSO control (Figure 3C). These compounds are in principle enhancing the efficacy of vaccinia in killing infected ID8 *Trp53^-/-^* cells.

**Figure 3.**
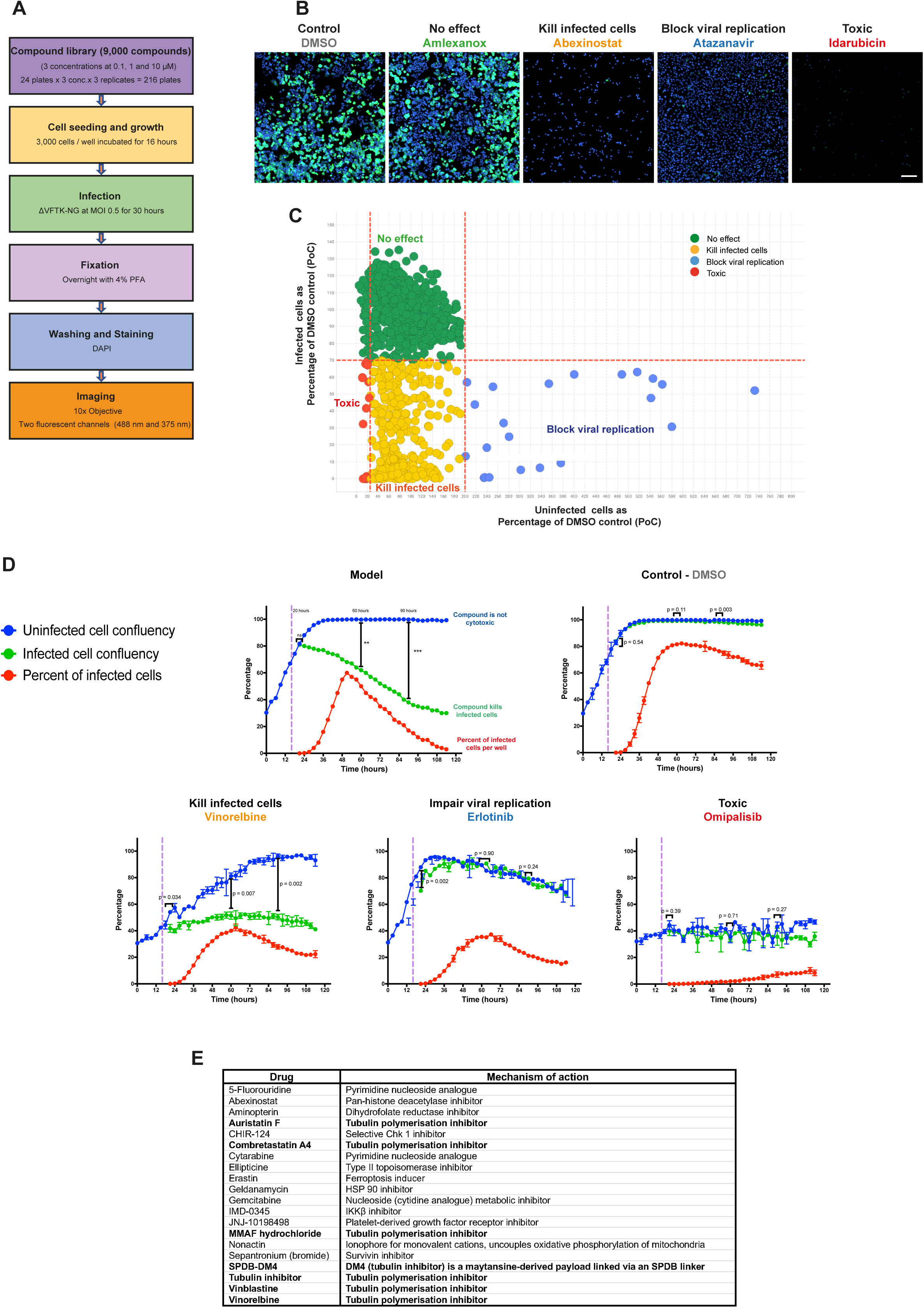
Identifying targets for combination therapies with oncolytic vaccinia. **A.** Schematic summarising the high throughput drug screening strategy. **B.** Representative images of ID8 *Trp53^-/-^*cells infected with ΔVFTK-NG virus and treated with the indicated compounds belonging to the four categories. NeonGreen expression (green) indicates infected cells and DAPI was used to stain nuclei (blue). Cells were imaged using Opera Phenix Plus with 10x air objective. Scale bar = 200 μm. **C.** Graphical illustration of compound allocation in the four categories. **D.** Representative graphs from the secondary validation screen of selected primary screen hits at 1μM (omipalisib, erlotinib and vinorelbine). The model represents an ideal compound and DMSO is the negative control. The purple dashed line represents the time at which ΔVFTK-NG virus (MOI 0.5) was added (16 hours post cell seeding). Statistical analysis was done by comparing uninfected against infected cell confluency at 20, 60 and 90 hours post cell seeding using Student’s t-test. Error bars represent standard deviation (SD). **E.** List of hit compounds after the secondary validation screen.

Images from ‘kill infected cells’ wells were reviewed and a cut-off ratio of infected to uninfected cells of 0.3 was applied to generate the initial hit list of 60 compounds (Table 1). During the validation of these hits, an additional 60 compounds, absent from the initial library that belong to the same class of molecules and/or had a similar mechanism of action were added to the hit list (Figure S2). A secondary live-cell imaging screen was conducted to assess compound cytotoxicity as well as the impact on viral replication and spread. Based on this analysis, the compounds were categorised as ‘kill infected cells’, ‘impair viral replication’ and ‘toxic’ (Figure 3D). Positive hits (‘kill infected cells’) were identified based on the reduction in median percentage of infected cell confluency per well over time compared to uninfected cells. This analysis resulted in twenty compounds, seven of which are tubulin polymerisation inhibitors (Figure 3E). We decided to focus on the seven compounds targeting the same process as this is likely to be indicative of true hits. The impact of these inhibitors was further confirmed in two additional human high grade serous ovarian cancer cell lines, OVCAR3 and OVCAR4 (Figure S3 and S4).

**Table 1.** List of 60 primary screen hits.

| Drug | Mechanism of Action |
| --- | --- |
| 2-amino-8-[4-(2-hydroxyethoxy) cyclohexyl]-6-(6-methoxy-3-pyridinyl)-4-methylpyrido[2,3-d]pyrimidin-7(8H)-one | PI3K/mTOR kinase inhibitor |
| 5-Fluorouridine | Pyrimidine nucleoside analogue |
| Abexinostat | Pan-histone deacetylase inhibitor |
| Actinomycin | DNA repair inhibitor |
| Alvespimycin (17-DMAG) hydrochloride | HSP 90 inhibitor |
| Aminopterin | Dihydrofolate reductase inhibitor |
| Amsacrine hydrochloride | Topoisomerase II inhibitor |
| AT13387 | HSP 90 inhibitor |
| Auristatin F | Tubulin polymerisation inhibitor |
| Bortezomib | Proteasome inhibitor |
| Canertinib | EGFR inhibitor |
| CCT137690 | Aurora kinase inhibitor |
| Cephaeline | Emetic alkaloid |
| CH5138303 | HSP 90 inhibitor |
| CHIR-124 | Selective Chk1 inhibitor |
| Chromomycin A3 | DNA replication and transcription inhibitor |
| Combretastatin A4 | Tubulin polymerisation inhibitor |
| Cytarabine | Pyrimidine nucleoside analogue |
| Cytochalasin E | Actin polymerisation inhibitor |
| D8-MMAD | Tubulin polymerisation inhibitor |
| D8-MMAF hydrochloride | Tubulin polymerisation inhibitor |
| Diacetoxyscirpenol | Ribosome inhibitor |
| Diosgenin | Precursor of steroidal hormones |
| Dp44mT | Iron chelator |
| Ellipticine | Topoisomerase II inhibitor |
| Emetine hydrochloride hydrate | Ribosome inhibitor |
| Enzastaurin | Protein kinase C inhibitor |
| Erastin | Ferroptosis inducer |
| Ergosterol | Sterol |
| Floxuridine | Pyrimidine nucleoside analogue |
| Fosbretabulin disodium | Tubulin polymerisation inhibitor |
| Ganetespib | HSP90 inhibitor |
| Geldanamycin | HSK90 inhibitor |
| Gemcitabine | Nucleoside (cytidine analogue) metabolic inhibitor |
| GSK3787 | PPARδ antagonist |
| HSP990 | HSP90 inhibitor |
| Imidazole ketone erastin | Ferroptosis inducer |
| MMAD | Tubulin polymerisation inhibitor |
| MMAF hydrochloride | Tubulin polymerisation inhibitor |
| MMAF-OMe | ADC cytotoxin and tubulin polymerisation inhibitor |
| NSC319726 | p53<sup>R175</sup> mutant reactivator |
| Nutlin-3 | P53-MDM2 interaction inhibitor |
| NVP-TAE226 | Focal adhesion kinase inhibitor |
| Pazopanib hydrochloride | Protein tyrosine kinase inhibitor |
| PDGF receptor tyrosine kinase inhibitor IV | Tyrosine kinase inhibitor |
| Premetrexed | Folate-dependent enzyme inhibitor. Inhibitor of nucleotide synthesis |
| Podophyllotoxin | Tubulin polymerisation inhibitor |
| Pralatrexate | Dihydrofolate reductase inhibitor |
| PU-H71 | HSP90 inhibitor |
| Raltitrexed | Thymidylate synthase inhibitor |
| Rosiglitazone | PPAR agonist |
| Silvestrol | RNA helicase inhibitor |
| Tanespimycin | HSP90 inhibitor |
| Thapsigargin | Sarco-endoplasmic reticulum Ca<sup>2+</sup> ATPase inhibitor |
| TRAM-34 | Potassium Channel inhibitor |
| Trifluridine | Nucleoside metabolic inhibitor |
| Tubulin inhibitor 1 | Tubulin polymerisation inhibitor |
| Vinblastine | Tubulin polymerisation inhibitor |
| Vincristine sulfate | Tubulin polymerisation inhibitor |
| Vinorelbine | Tubulin polymerisation inhibitor |

Vinorelbine is a semi-synthetic second generation vinca alkaloid with a broad spectrum antitumour activity that inhibits microtubule assembly.^31,32^ It has reduced side effects including neurotoxicity compared to other vinca alkaloids.^33^ An oral vinorelbine formulation is also available making it an appealing combination agent.^34^ Vinorelbine is mainly used to treat non-small cell lung and breast cancers but also has some efficacy against ovarian cancer.^35,36,37,38^ Based on this, we selected vinorelbine for mechanistic exploration and *in vivo* experiments.

### Vaccinia and vinorelbine induce ID8 *Trp53^-/-^* cell death via apoptosis

Vinorelbine treatment of uninfected ID8 *Trp53^-/-^* cells for 16 hours results in the loss of their characteristic cobblestone appearance and cell to cell contacts as well as some cell rounding, indicative of possible progression to cell death (Figure 4A). Immunofluorescence analysis confirmed that vinorelbine results in loss of microtubules in both ID8 *Trp53*^-/-^ cells with or without infection with the ΔVFTK virus (Figure 4B). Vinorelbine treatment also induced nuclear fragmentation and the formation of multinucleated cells suggestive of defects in cytokinesis or cell fusion.

**Figure 4.**
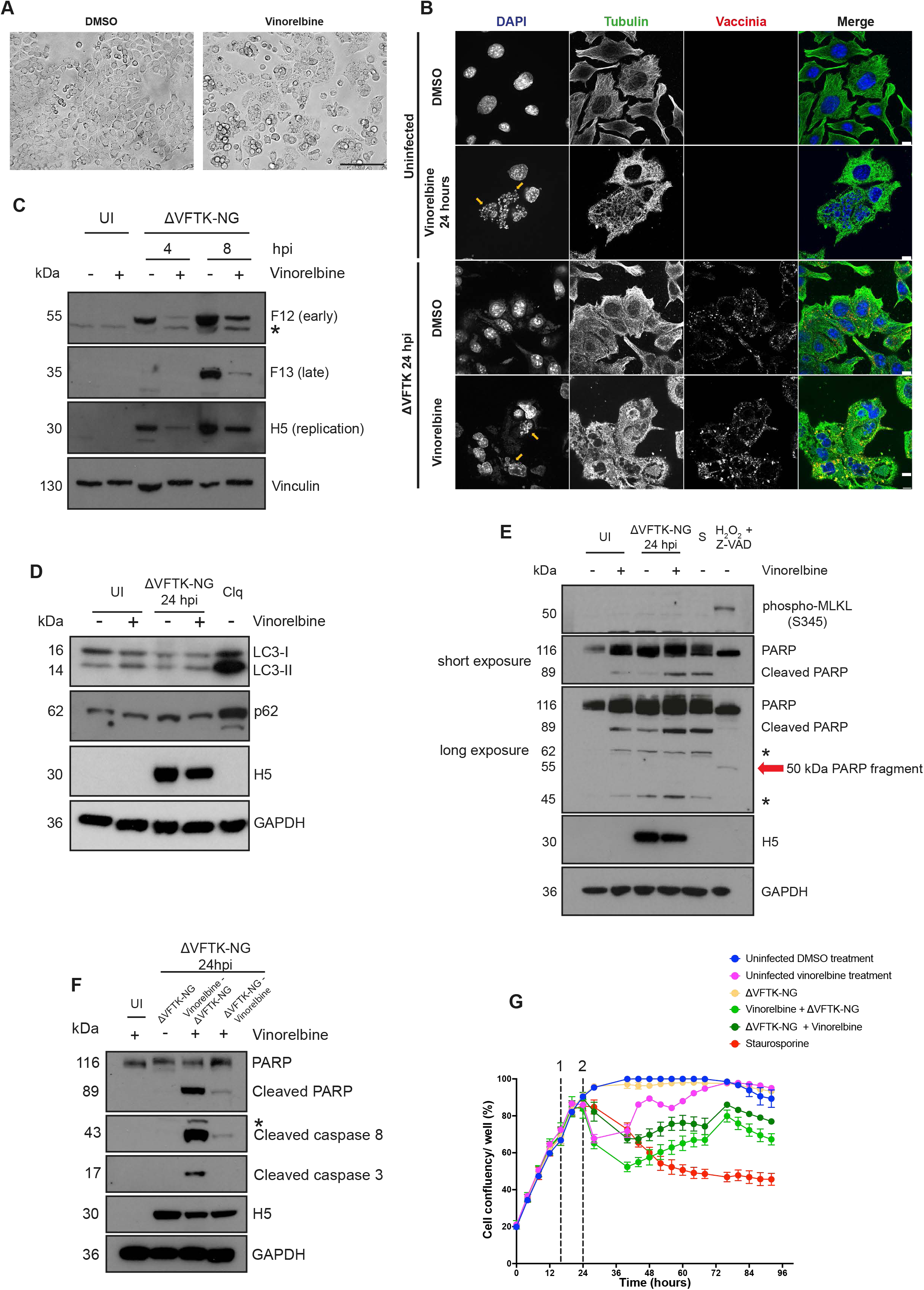
Vaccinia and vinorelbine induces apoptotic ID8 *Trp53^-/-^* cell death. **A.** Representative phase contrast images of ID8 *Trp53^-/-^* cells after 16 hours treatment with DMSO or vinorelbine (1 μM). Scale bar = 150 μm. **B.** Immunofluorescent images of microtubule cytoskeleton (Tubulin - green) in ID8 *Trp53^-/-^* cells treated with DMSO or vinorelbine with or without infection with ΔVFTK (Vaccinia - red). DAPI (blue) was used to stain nuclei and cytoplasmic viral factories. Yellow arrows indicate fragmented nuclei. Scale bar = 10 μM. **C.** Immunoblot of the indicated viral proteins following infection for 4 or 8 hours with ΔVFTK-NG following 16 hours pre-treatment with DMSO or vinorelbine. Vinculin is the cell loading control. **D.** Immunoblot examining the levels of LC3 lipidation (LC3-II) and p62 in ID8 *Tr53^-/-^* cells pretreated with vinorelbine or DMSO for 8 hours and infected with ΔVFTK-NG for 24 hours. Chloroquine (Clq) treatment for 3 hours represents the positive control. **E.** Immunoblot examining the levels pf MLKL phosphorylation and PARP cleavage in ID8 *Tr53^-/-^* cells pretreated with vinorelbine or DMSO for 8 hours and infected with ΔVFTK-NG for 24 hours. Staurosporine (S) (8 hours) and 0.1% hydrogen peroxide (H_2_O_2_) combined with Z-VAD (two hours) represent positive controls for apoptosis and necroptosis respectively. **F.** Immunoblot examining the levels of cleaved caspase 8, 3 and PARP in ID8 *Tr53^-/-^* cells treated with vinorelbine 8 hours before or after infection with ΔVFTK-NG for 24 hours. In D, E and F H5 and GAPDH represent the viral and cell loading controls, respectively. For all experiments, uninfected DMSO-treated cells (UI) were the negative control. Asterisks (*) indicate non-specific bands. Immunoblot experiments were repeated three times and representative examples are shown. **G.** Median percentage cell confluency over time for the indicated conditions. Cells were treated with the indicated compounds or ΔVFTK-NG at 16 hours post seeding (dashed line numbered 1). The dashed line numbered 2 (24 hours post cell seeding) represents the time of addition of either vinorelbine or ΔVFTK-NG for the combination groups.

Vaccinia hijacks the microtubule cytoskeleton to facilitate its replication, intracellular transport and spread.^39,40,41,42^ We previously found that depolymerization of microtubules with nocodazole reduces virus yield.^43^ Consistent with this, immunoblot analysis revealed that vinorelbine treatment reduced both early (F12) and late (F13) viral protein expression, as well as H5, which is required for viral replication (Figure 4C). Vinorelbine treatment prior to infection of ID8 *Trp53^-/-^* cells clearly impairs viral gene expression but it is not immediately obvious why there is enhanced cell death or which programmed cell death pathway (autophagy, necroptosis or apoptosis) is activated.

To assess whether vinorelbine induces autophagy in ID8 *Trp53^-/-^* infected with the ΔVFTK-NG virus, we performed immunoblot analysis to examine the level of p62 expression and LC3 lipidation (LC3-II) (Figure 4D). In contrast to chloroquine treated cells (positive control), there was no change in the level of p62 in infected or non-infected cells with or without vinorelbine. Likewise, the levels of LC3-II did not change, except in the positive control indicating that autophagy is not responsible for increasing cell death of vinorelbine treated ID8 *Trp53^-/-^* cells.

To investigate the possible involvement of necroptosis, we performed immunoblot analysis to examine the level of MLKL phosphorylation (pMLKL) and look for the presence of 50 kDa PARP fragment generated by lysosomal proteases.^44^ We found pMLKL and the 50 kDa PARP fragment which are indicative of necroptosis are only present in the positive control (0.1% hydrogen peroxide and Z-VAD treatment) (Figure 4E). We did, however, see that vinorelbine induced cleavage of PARP to generate an 89 kDa fragment in infected cells, suggesting that apoptosis is responsible for their death.^45^ The presence of cleaved caspase 3 and 8 in immunoblots of infected cells treated with vinorelbine confirmed apoptosis was activated (Figure 4F). The combination of vinorelbine and infection is required to induce apoptosis as PARP, caspase 3 or 8 cleavage is not observed in non-infected or infected cells with or without the drug respectively. Pre-treating infected cells with vinorelbine also appears to be more effective at inducing PARP, caspase 3 or 8 cleavage (Figure 4F). To examine if this translates into increased cell death, we performed live cell imaging of ID8 *Trp53^-/-^* cells treated with vinorelbine before or after vaccinia infection using the same conditions used in the initial drug screen. We found that pre-treating ID8 *Trp53^-/-^* cells with vinorelbine leads to increased initial cell death compared to adding vinorelbine after infection (Figure 4G). However, both conditions eventually result in similar levels of cell death.

### Vinorelbine and ΔVFTK-NG-GM-CSF improves mouse survival

An *in vivo* distribution study was carried out to examine the tumour specificity of the ΔVFTK-NG-GM-CSF virus (Figure 5A). Viral replication was detected in omental tumours but not in liver or spleen in mice inoculated intraperitoneally with virus with or without vinorelbine (Figures 5B and C). Pre-treatment with vinorelbine impacted viral replication *in vivo* consistent with our observations in ID8 *Trp53^-/-^* cells in culture (Figures 5B and D). Vinorelbine also reduced the level of GM-CSF expression in omental tumours (Figure S5). To assess the efficacy of the combination treatment on the survival of mice we treated mice with vinorelbine 24 hours before injection of the ΔVFTK-NG-GM-CSF virus (Figure 5E). This combination prolonged mouse survival compared to vinorelbine (median survival of 73.5 compared to 69 days, p = 0.022) (Figure 5F). Moreover, both conditions were also significantly better than virus alone (median 57 days). Our observations suggest that combining vinorelbine and ΔVFTK-NG-GM-CSF virus improves mouse survival against high grade serous ovarian carcinomas.

**Figure 5.**
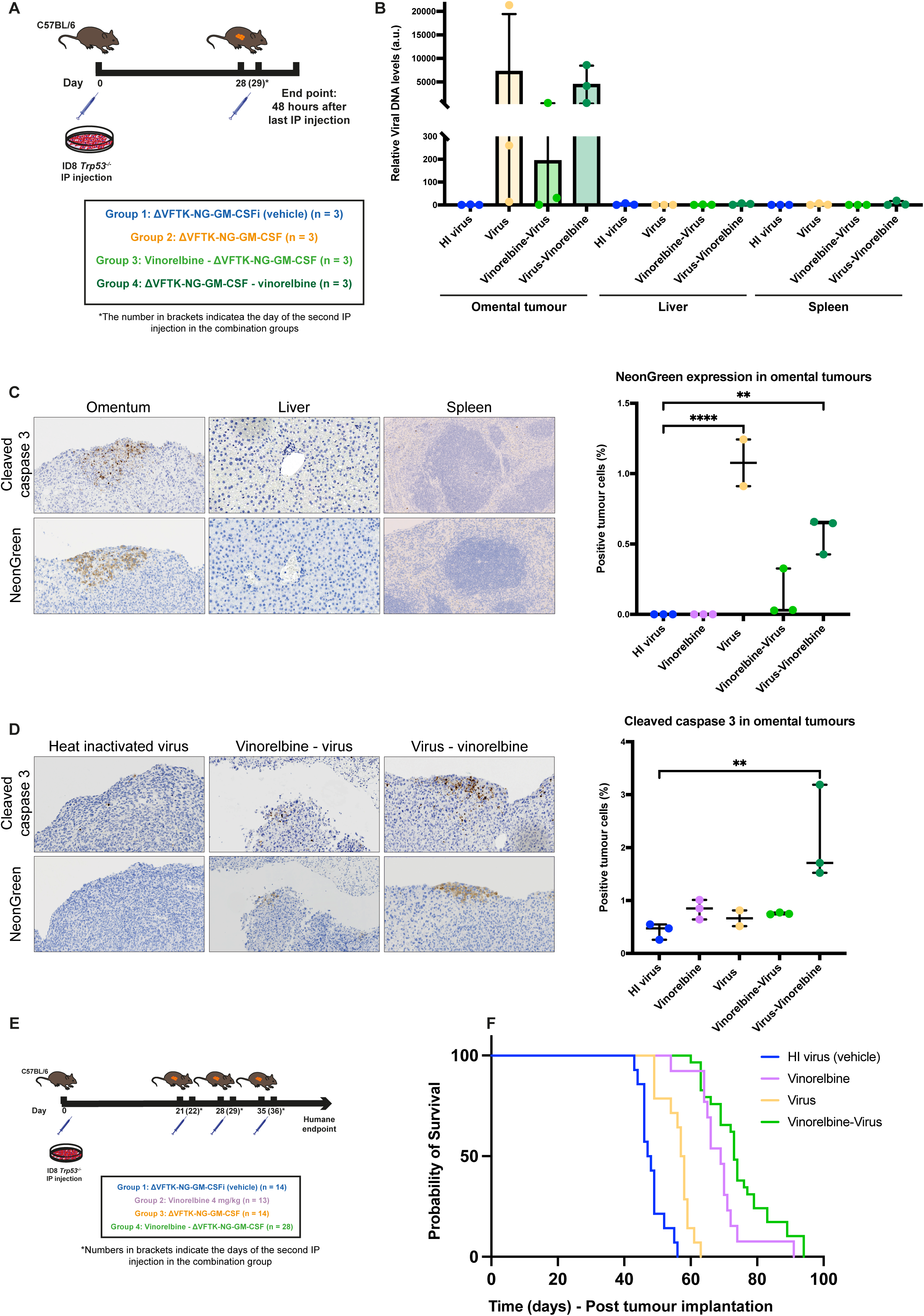
Assessing the efficacy of ΔVFTK-NG-GM-CSF in vivo. **A.** Schematic representation of the experimental design of the distribution study. Mice were injected IP with ID8 *Trp53^-/-^*cells on day 0 and received their IP injection on day 28 and the combination groups received their second component (vinorelbine or virus) on day 29. Heat inactivated (HI) virus (ΔVFTK-NG-GM-CSFi) is the negative control. **B.** Quantification of viral DNA in omental tumour, liver and spleen measured by qPCR and expressed relative to viral DNA levels of the liver sample in the HI virus group. Error bars represent mean ± SD. **C.** Representative immunohistochemical images of the distribution of cleaved caspase 3 and NeonGreen in omental tumour, liver and spleen harvested from a mouse in Group 4. The graph shows the quantification of NeonGreen positive cells in omental tumours. **D.** Representative immunohistochemical images of the distribution of cleaved caspase 3 and NeonGreen in omental tumours from mice in Groups 1, 3 and 4. The graph shows the quantification of cleaved caspase 3 positive cells in omental tumours. **E.** Schematic representation of the experimental design of the vaccinia-vinorelbine combination study. Mice were injected with ID8 *Trp53^-/-^* cells on day 0 and started receiving their IP treatment injections on day 21. Mice allocated to single treatment groups received their inoculations on days 21, 28 and 35 and those allocated to the combination groups received vinorelbine and ΔVFTK-NG-GM-CSF 24 hours apart. Heat inactivated (HI) ΔVFTK-NG-GM-CSFi virus was the negative control. **F.** Kaplan-Meier survival curve showing survival data for each group analysed by log-rank test. One mouse belonging to group 2 was excluded from the analysis as after IP injection of ID8 *Trp53 ^-/-^*cells omental tumour failed to form. The analysis is from the combination of two survival experiments following the same protocols.

## Discussion

Vaccinia virus has the promise of being an ideal oncolytic agent given its safety profile and ability to be genetically enhanced as well as combined with drug treatments.^46,47^ Clinical trials frequently use viruses deleted of genes encoding VGF (ΔVGF) and/or the thymidine kinase (ΔTK) as this increases their tumour specificity and safety.^48,17^ Deletion of F1, an inhibitor of apoptosis in combination with TK also improves the oncolytic effectiveness of the virus as it extends mouse survival in a human glioblastoma model and delays tumour growth in a syngeneic mouse colon model.^26,27,49^ Based on the available evidence, a virus lacking VGF, TK and F1 is predicted to be safer and induce more tumour cell death.

We have now deleted VGF, TK and F1 in the Western reserve (WR) strain of Vaccinia and also inserted GM-CSF to the genome to increase the immunogenicity of the virus. NeonGreen was also added which like other reporters, can be used to monitor viral replication, distribution, detect metastatic disease and assess toxicity clinically.^50,51^ Our *in vivo* survival studies demonstrated that our new recombinant viruses were well tolerated in mice bearing ID8 *Trp53^-/-^* omental tumours but that only the ΔVFTK-NG-GM-CSF virus had consistently significant survival benefit (Figure 2B and S1). Moreover, when injected intraperitoneally the ΔVFTK-NG-GM-CSF virus specifically replicates in omental tumours but not in the liver or spleen (Figure 5A-C).

Multiple early phase clinical trials demonstrate that even with genomic modifications that enhance its tumour specificity, oncolytic activity and immunogenicity, vaccinia clearly still needs a combination agent to achieve its full oncolytic potential.^25,46^ Given this, we performed a high throughput drug screen on ID8 *Trp53^-/-^*cells using 9,000 well-characterised compounds to identify those that increased the killing efficiency of our triple deleted recombinant virus. By performing primary and secondary drug screens, we identified 20 hit compounds, seven of which were tubulin polymerisation inhibitors (Figure 3E). Vinorelbine was our drug screen hit compound of choice given it has some activity against ovarian cancer.^35,36,37^

We found that the combination of vaccinia with vinorelbine induces ID8 *Trp53^-/-^* cell death via apoptosis. This is not unexpected, as the ΔVFTK-NG virus lacks F1 the main inhibitor of intrinsic apoptosis during infection.^26,27^ Vinca alkaloids such as vinorelbine also induce apoptosis, which is often attributed to impaired spindle formation and subsequent prolonged mitotic arrest.^52,53^ However, these chemotherapeutics can additionally stimulate apoptosis independently of the cell cycle via alternative mechanisms including the activation of intrinsic apoptosis.^54^ Microtubule targeting agents have been shown to upregulate pro-apoptotic Bcl-2 proteins (BAX, BAK, PUMA, Noxa and Bad) as well as inactivate anti-apoptotic Bcl-2 proteins through phosphorylation (Bcl-2, Bcl-xL, Mcl-1, Bcl-W).^53,55,56^ Vinorelbine and the ΔVFTK-NG virus both act on the intrinsic mitochondrial apoptotic pathway and their effects complement each other leading to enhanced apoptotic cell death compared to either agent alone. Therefore, vinorelbine pre-treatment will not only induce mitotic arrest but also activate BAK/BAX leading to mitochondrial membrane pore formation resulting in a pro-apoptotic ‘priming’ making cells more susceptible to vaccinia induced apoptosis.^57^ Consistent with this notion, vinorelbine pre-treatment of cells resulted in increased PARP and caspase cleavage at 24 hours post infection compared to adding the drug after infection (Figure 4F). Live-cell imaging also demonstrated that addition of vinorelbine before infection increased initial cell death compared to first adding the virus and then the drug (Figure 4G).

When tested in a mouse model of high grade serous ovarian carcinoma, we found that the combination of vinorelbine followed by infection with the ΔVFTK-NG-GM-CSF virus produced a significant survival benefit compared to the control (median survival of 73.5 compared to 47.5 days). Our results are in line with previous studies demonstrating that a combination of vinorelbine and vesicular stomatitis virus (VSV) significantly prolonged the survival of aggressive subcutaneous 4T1 syngeneic mouse model of triple negative breast cancer.^58^ Vinorelbine has good response rates in phase I and II clinical trials for recurrent ovarian cancer and the National Comprehensive Cancer Network recommended vinorelbine for recurrent epithelial, fallopian tube or primary peritoneal cancers.^59^ It is evident from our *in vivo* experiments that vinorelbine at the dose used (4mg/kg) has good efficacy against ID8 *Trp53-/-* tumours, suggesting that the drug represents a promising therapy for patients with high grade serous carcinoma. Nevertheless, when vinorelbine is combined with the ΔVFTK-NG-GM-CSF virus mouse survival is further increased.

Our findings suggest that the enhanced tumour cell killing observed with the combination of vinorelbine and ΔVFTK-NG-GM-CSF is primarily due to an additive or synthetic lethal interaction, rather than increased viral replication. Although vinorelbine initially suppressed cell proliferation in vitro, confluency recovered over time, indicating a predominantly cytostatic effect (Figure 4G). In contrast, the combination treatment led to a sustained reduction in confluency, consistent with induction of apoptosis as confirmed by cleaved caspase-3 and PARP immunoblotting (Figure 4F and G). Moreover, qPCR analysis of viral gene expression demonstrated that vinorelbine pre-treatment reduced viral replication relative to infection with virus followed by vinorelbine, demonstrating that vinorelbine impairs vaccinia replication when administered prior to viral infection. These data support the notion that the combination effect arises from enhanced cell death rather than increased viral propagation.

Further experiments are required to examine the impact on mouse survival of reducing the number of doses and concentration of vinorelbine together with higher viral titres. In this way it should be possible to develop strategies with the ΔVFTK-NG-GM-CSF virus that are effective with lower vinorelbine doses to decrease systemic toxicity and the likelihood of patients becoming resistant to vinorelbine. It will also be important to examine the immunostimulatory function of GM-CSF in our combination approach. Future studies will also focus on validating these findings across additional syngeneic and human ovarian cancer models to assess the robustness and generalizability of the therapeutic effect.

## Materials and methods

### Cells and culture

The ID8 *Trp53^-/-^* cell line was provided by Professor Iain McNeish, Imperial College London (London, UK).^60^ HeLa and BS-C-1 cell lines were provided by the European Molecular Biology Laboratory (Heidelberg, Germany) and 143B TK^-/-^ from The Francis Crick Institute Cell Services (London, UK). All cell lines were mycoplasma tested and maintained in Dulbecco’s Modified Eagle’s Medium (DMEM) (Sigma-Aldrich, St. Louis, Missouri, #51435C) supplemented with 10% fetal bovine serum (FBS) (Thermo Fisher Scientific, Waltham, Massachusetts, #10270-106) and 1% penicillin/ streptomycin (pen/strep) (Sigma-Aldrich, #P4333) at 37°C and 5% CO_2_.

### Virus amplification and sucrose purification

HeLa cells were grown in 15 cm culture dishes (Corning Inc., Sigma Aldrich, #353025) at approximately 80% confluency and were infected with vaccinia at a MOI of 0.1 for 48-72 hours. The cells were scraped in PBS and centrifuged at 500 x g at 4°C for five minutes. Cell pellets were resuspended in PBS and re-centrifuged. The resulting cell pellet was resuspended in 7 ml of Tris buffer (10 mM Tris HCl, 2 mM MgCl_2_, pH 9.0), disrupted by 20 strokes in a 7 ml Wheaton Dounce homogenizer (DWK Life Sciences, Millville, New Jersey, #357538) and centrifuged at 500 x g at 4°C for five minutes. The supernatant was collected and added on top of an 8 ml 35% sucrose cushion in a Beckman SW40 ultracentrifuge tube (Beckman Coulter, Brea, California #3117-0380). The solution was centrifuged at 192, 000 x g at 4°C for 30 minutes in a Beckman Optima l-100 XP ultracentrifuge, using an SW32Ti swing-out rotor. The virus pellet was resuspended in a desired amount of Tris buffer, titrated using plaque assays, aliquoted and stored at -20°C (short term) or -80°C (long term).

### Viral plaque assays and heat inactivation

Confluent monolayers of BS-C-1 cells were infected with vaccinia virus in serum free media at the required dilutions for one hour at 37°C, 5% CO_2_. The media was then replaced with 2 ml of semi-solid media (1:1 ratio of 3% Carboxy-methyl cellulose sodium salt dissolved in water and 2X Modified Eagle’s Medium (MEM) supplemented with 10% FBS and 1% pen/ strep). After 72 hours, the cells were fixed by adding 1 ml of 8% formaldehyde to the media for 30 minutes at room temperature. The media and formaldehyde were removed, and cells were stained with 1 ml crystal violet (1:5 dilution in Phosphate buffered saline [PBS]) for 30 minutes. Subsequently, the plates were rinsed with cold water and left to air dry. To determine the plaque size, plaque diameter was measured in mm using Fiji line tool, available at imagej.net. For heat inactivation, aliquots of viruses were incubated for three hours at 60°C and then cooled on ice for one hour. Heat inactivation was confirmed by the absence of plaque forming units using plaque assays.

### Generation of recombinant viruses

Recombinant viruses were constructed in the ΔVF virus background which is derived from the Western Reserve (WR) strain.^30^ The transient dominant selection (TDS) method was used to delete TK from the ΔVF genome.^61,62^ To create the targeting vector for TDS a DNA fragment containing the left (387bp) and right (321bp) TK recombination arms was cloned into the pSSGB vector containing the selectable markers GFP-*bsd* under the control of a synthetic vaccinia promoter (pSS) . ^62^ The fluorescent ΔVFTK virus expressing NG (ΔVFTK-NG) and the ΔVFTK virus expressing NG and GM-CSF (ΔVFTK-NG-GM-CSF) under the control of the synthetic early/late vaccinia promoter (pEL) at the TK locus were created via homologous recombination between ΔVF virus and the pBSΔTK-NG and pBSΔTK-NG-GM-CSF targeting vectors, respectively. These targeting vectors were created using BlueScript (PBS(+)) as the backbone vector and DNA fragments containing sequences for pEL-NG and pEL-NG-GM-CSF as the inserts. For all new recombinant viruses, the targeting vectors contained TK recombination arms flanking the DNA sequence to be inserted in the vaccinia genome. A small portion of the 5’ open reading frame of J2R (gene encoding for TK) was retained within the left TK recombination arm to ensure the correct transcription of J1R. These DNA fragments were synthesised using the IDT gBlocks gene synthesis service (Coralville, Iowa). Subsequently, HeLa cells, infected with ΔVF at MOI of 0.05 were transfected with the relevant targeting vectors using Lipofectamine 2000 (Thermo Fisher Scientific, #52887) and incubated for 48 hours at 37°C and 5% CO_2_. The viruses were harvested by scraping the cells and subjecting them to three freeze-thaw cycles. The harvested viruses were used to infect confluent monolayers of 143B TK^-/-^ cells with BrdU selection or BS-C-1 cells with or without blasticidin selection. Recombinant viruses, which are shown schematically in Figure S6, were isolated by identifying and picking fluorescent green plaques over at least three rounds of purification. For all three recombinant viruses, TK deletion and/or insertions of NG or NG-GM-CSF were confirmed by PCR analysis after each round of plaque purification and sequencing. The recombinant viruses were amplified and purified using a 35% sucrose gradient.

### Quantification of viral DNA and GM-CSF mRNA

Total DNA was isolated from mouse tissues using DNeasy Blood and Tissue Kit (QIAGEN, Hilden, Germany), as per manufacturer’s protocol and quantified using a NanoDrop (Thermo Fisher Scientific). Real-time quantitative PCR (qPCR) was performed in triplicates in 384-well reaction plates (Applied Biosystems, Waltham, Massachusetts). Each PCR reaction (total volume 10 μl) contained 20 ng DNA, 5 μl Power SYBR-Green PCR Master Mix (Applied Biosystems) and 0.5 μM of forward and reverse primers. Viral DNA was amplified using primers specific to a region for H5 vaccinia gene (H5-F: 5’-GTAAGAAGTAAATGCGTGC-3’, H5-R: 5’-CCACGTTTGTTCATATACTAC-3’) and APP1 (APP1-F: 5’-CGGAAACGACGCTCTCATG-3’ and APP1-R: 5’-CCAGGCTGAATTCCCCAT-3) was amplified as a nuclear gene standard reference. Changes in viral DNA amount were calculated using the 2^-ΔΔCt^ method and represented as fold changes relative to the indicated control.^63^ For quantification of GM-CSF mRNA, 20 mg of omental tumour tissue was homogenised in TRIzol reagent using a Precellys Evolution tissue homogenizer (Bertin Technologies, Montigny-le-Bretonneux, France). Following chloroform addition and centrifugation, the aqueous phase containing RNA was combined with an equal volume of 70% ethanol and purified using RNeasy Mini Kit columns, according to the manufacturer’s instructions (QIAGEN). One-step quantitative RT-PCR was then performed on 50 μg of RNA template using the QuantiNova® SYBR® Green RT-PCR Kit (QIAGEN) with 0.5 μM of gene-specific primers: GM-CSF (Csf2-F: CTACTACCAGACATACTGCC; Csf2-R: GCATTCAAAGGGGATATCAG) and GAPDH as the housekeeping control (GAPDH-F: TCTTGTGCAGTGCCAGCCT; GAPDH-R: CAATATGGCCAAATCCGTTCA).

### Immunoblotting

Cells were collected and lysed in 21 μl of PBS supplemented with Protease/ Phosphatase inhibitors (Cell Signalling, #5871) and 5U benzonase nuclease (Millipore, #E1014). SDS was added at a final concentration of 1% followed by 22 μl of 2X SDS loading buffer (Thermo Fisher Scientific, #LC2676). The samples were heated at 95°C for three minutes, loaded onto Bolt 4-12% or 10% Bis-Tris Plus pre-cast gels (Thermo Fisher Scientific) and run in MOPS SDS running buffer (Thermo Fischer Scientific, #NP0001) for 55 minutes at 150 V. SeeBluePlus2 protein standard (Thermo Fisher Scientific, #LC5925) was used as reference for protein molecular weight. The proteins were transferred to nitrocellulose membranes (Thermo Fischer Scientific, #IB23001), blocked in 5% milk in PBS with 0.1% Tween20 (PBS-T) (Sigma-Aldrich, #P9416) for one hour at room temperature and incubated overnight with primary antibodies. Primary antibodies used were: F12 (1:4,000, ^64^), F13 (1:6,000, ^39^), H5 (1:10,000, ^65^), GRB2 (1:1,000, Santa Cruz, Dallas, Texas, #sc-255), Vinculin (1:10,000, Sigma-Aldrich, #V9264), GAPDH (1:1,000, Santa Cruz, #sc-32233), PARP (1:1,000, Cell Signalling, Danvers, Massachusetts, #9542), cleaved caspase 8 (1:1,000, Cell Signalling, #8592), cleaved caspase 3 (1:1,000, Cell Signalling, #9664), LC3-B (1:1,000, Abcam [#ab48394], Cambridge, UK), p62/SQSTM1 (1:1,000, Novus Biologicals, Centennial, Colorado, #NBP1-42822) and NeonGreen (1:1,000, Cell Signalling, #41236). Goat anti-rabbit (#111035003) and anti-mouse (#115005003) secondary HRP antibodies, obtained from Jackson ImmunoResearch (West Grove, Pennsylvania), were used at 1:10,000 in 5% milk with PBS-T. For the examination of phosphorylated proteins, 1 mM orthovanadate (New England Biolabs, Ipswich, Massachusetts, #P0758) was added during blocking, primary and secondary antibody incubation. The membranes were incubated with SuperSignal West Pico PLUS Chemiluminescence reagent (Thermo Fischer Scientific, #34580) for one minute at room temperature and exposed to UltraCruz Autordiography Film (Santa Cruz, #sc-201697).

### Drug screen Chemical library

A small molecule drug library of 9,000 well-characterised compounds, assembled from commercial libraries by the High Throughput Screening (HTS) science technology platform (STP) of the Francis Crick Institute was used for the primary screen. This chemical library can be accessed (https://hts.crick.ac.uk/db/view/libraryView.php) under the database name: ‘Full Chemical Collection V6’. A customised library of 120 compounds was purchased form MedChemExpress (Monmouth Junction, New Jersey) for the secondary screen.

### Primary screen

The small molecule library, resuspended in dimethyl sulfoxide (DMSO), was transferred into intermediate Low Dead Volume (LDV; Labcyte, #LPS-0200) 384-well plates at a concentration of 10 and 1 mM. An acoustic liquid handler (Echo 550 Beckman-Coulter) was then used to dispense compounds into Greiner microclear 384-well plates (Greiner, #781091) so they reach a final concentration of 0.1, 1 and 10 μM. Each assay plate was prepared in triplicate. Prior to cell seeding, the compounds were diluted in 10 μl of complete DMEM per well. A volume of 40 μl of cells, corresponding to 3,000 cells, was dispensed into the 384-well assay plates, which were incubated for 16 hours at 5% CO_2_ and 37°C. The ΔVFTK-NG virus was dispensed in each well at an MOI 0.5 and the plates were incubated at 5% CO_2_ and 37°C for 30 hours. Cells were fixed using 4% paraformaldehyde (PFA), permeabilised and stained by adding 0.01% Triton X-100 and DAPI. The cells were imaged on Opera Phenix^®^ Plus (Revvity, Waltham, Massachusetts) using 10x air NA 0.3 objective. The microscope was equipped with a set of lasers and filters for the excitation and emission wavelengths specific for DAPI (excitation: 375 nm and emission: 435-480 nm) and NeonGreen (excitation: 488 nm and emission: 500-550 nm).

### Primary drug screen analysis

Five fields per well were imaged and analysed using Harmony^®^ (version 5.0). Screen data were analysed with the cellHTS2 R package.^66,67^ Raw measurements were scaled relative to the mean of the within-plate DMSO controls to report a percentage-of-control (PoC) for each feature and replicates were summarised as a median value. The replication between technical replicates was assessed using the Spearman correlation. Tibco Spotfire (version 14.0) software was used to process the large datasets. Hit determination was based on how the compounds compared against PoC in terms of infected and uninfected cells.

### Animal experiments

All animal experiments were carried out in accordance with the UK Home Office regulations under the project licence PA780D61A and PP1321516 (from May 2024) and were approved by the Imperial College Animal Welfare and Ethical Review Body. Female, six-to seven-week-old C57BL/6 mice were purchased from Charles River Laboratories (Harlow, UK) and acclimatised for a week prior to cell injection. They had standard laboratory diet and free access to water. Mice were inoculated intraperitoneally (IP) with ID8 *Trp53^-/-^* cells (5×10^6^) in 200 μl sterile PBS on day 0. The tumours were allowed to grow for 14, 21 or 28 days depending on the protocol of each study. Details of the experimental designs of these studies are provided in Figures 2, 5 and S1. Day 21 was used as the standard tumour establishment time point used in most *in vivo* survival studies, as it reflects late-stage disease. In the survival study presented in Figure S1, viral treatment began on day 14, and mice received four intraperitoneal (IP) viral injections instead of three. This modified protocol was designed to assess whether initiating treatment earlier, when tumour burden is lower, and increasing the number of viral doses could enhance therapeutic efficacy. Day 28 was used exclusively for the viral distribution study. This later time point was chosen to allow for greater tumour progression, providing larger tissue samples suitable for assessing viral localisation within the tumour. Sample sizes were established with the help of the experimental design assistant of NC3Rs to achieve statistical power while minimising animal use. All viruses were injected IP at a titre of 1×10^7^ pfu/ml in 200 μl sterile PBS. Vinorelbine tartrate was obtained from Hammersmith hospital pharmacy, Imperial Collage Healthcare NHS Trust (London, UK) and was injected IP at 4 mg/kg in 200 μl sterile PBS. The mice were weighed daily, monitored regularly for adverse side effects, and were killed when reached humane endpoints. These included any of the following: swelling restricting movement, piloerection, hunched posture, reduced activity, facial grimace, dehydration lasting > 24 hours, 15% weight loss, altered respiration and self-mutilation. All decisions on animal welfare were made by staff blinded to treatment allocation. The schedule 1 method followed was cervical dislocation and exsanguination by decapitation. Omental tumours, liver and spleen were dissected and placed in 10% neutral buffered formalin for 48 hours and then transferred to 70% ethanol.

### Histopathology

Harvested tissues, embedded in paraffin to create blocks, were sectioned at 3 μm thickness and attached on glass microscope slides for examination. Histochemical Haematoxylin and Eosin (H&E) staining was carried out using Tissue-Tek Prisma^®^ *Plus* Automated Slide Stainer (Leica Biosystems, Deer Park, Illinois). Blinded histopathological analysis was performed by Professor Priestnall and Dr Suarez-Bonnet, board-certified veterinary pathologists associated with the Experimental Histopathology STP of The Francis Crick Institute. For immunohistochemistry (IHC), samples were stained for NeonGreen (1:200, Cell Signalling, #41236) and cleaved caspase 3 (1:250, Cell signalling, #9579S) antibodies, respectively. IHC staining was performed on the Leica Bond Rx autostainer platform (Leica Biosystems, #3498240) and a DAB kit which included a secondary antibody, DAB and haematoxylin counterstain was used (BOND Polymer Refine Detection, Leica Biosystems, #DS9800). The slides were scanned with a Zeiss Axio Scan Z1 Slide Scanner operated by ZEN Lite Software.

Quantitative analysis of immunohistochemistry slides was performed using QuPath (v0.3.0). Slides were scanned and imported from a secure server. Cell detection was carried out using default DAB channel parameters, matched to the chromogen used. Stain separation was optimised by estimating stain vectors. Tumour and non-tumour regions were annotated to train a random forest classifier for compartment classification. Cleaved caspase 3 and vaccinia staining were analysed using intensity thresholds of 0.15 and 0.2 respectively (Cell: DAB OD Mean). A script was developed to calculate the percentage of marker expression in tumour versus non-tumour areas, with results reported as intra-tumoral expression percentages.

### Immunofluorescence

For fixed cell imaging, cells were seeded on 0.1% fibronectin-coated (Sigma-Aldrich, #F0895) coverslips. Cells were infected with vaccinia virus and at experimental endpoint, cells were fixed with 1ml of iced cold methanol for 20 minutes at -20°C and then washed with PBS. Cells were incubated in blocking buffer (1% bovine serum albumin and 2% fetal calf serum) for 30 minutes followed by the primary antibody for one hour. Primary antibodies used were as follows: B5 (1:1,000 ^68^) and ɑ - Tubulin (1:500, Sigma-Aldrich, #T6074). Cells were incubated in Alexa Fluor 488 and Alexa Fluor 568 conjugated secondary antibodies (1:1,000 dilution in blocking buffer) for a further 40 minutes and stained with DAPI (300 nM in PBS) for five minutes. Coverslips were mounted on glass microscope slides using 5 μl Mowiol. Mounted coverslips were imaged on a Zeis Axio Observer spinning-disk microscope equipped with a Plan-Apochromat 100x/1.46 oil lens, an Evolve 512 cameral and a Yokagawa CSUX spinning disk. The microscope was controlled by Slidebook software (3i Intelligent Imaging Innovations). Images were analysed using Fiji Imaging Analysis Software.

### Live-cell imaging

ID8 *Trp53^-/-^* cells were seeded at the desired confluency in 384-well plates and were transferred to the Incucyte S3 live-cell imaging system (Sartorius) to capture phase-contrast and green-fluorescent images every three hours using a 4x objective. Cells were removed from the Incucyte S3 at the required timepoints for drug treatment and/or infection and quickly replaced in the Incucyte to continue imaging. Whole wells were imaged and analysed using the integrated software module “Basic Analyser”. The median percentage confluency of infected and uninfected cells and the ratio of green fluorescence per well area were quantified. Raw data was exported for further statistical analysis.

Live cells to assess morphological changes following vinorelbine treatment were imaged using Invitrogen Evos^TM^ M5000 microscope (Invitrogen) equipped with advanced LED illumination and a range of objective lenses to achieve various magnifications. Images were captured using the integrated software.

## Statistical analysis

Statistical analysis was performed using Prism 10 (GraphPad Software). In all graphs, data are represented by the mean and standard deviations (SD) or median and interquartile range (IQR) from three independent experiments, unless stated otherwise. Data were tested for normality of distribution using normality and lognormality tests in Prism 10. For parametric data: Student’s t test was used to compare two data sets and one-way ANOVA for multiple data sets followed by either a Tukey’s (comparing samples to each other) or Dunnett’s (comparing samples to control) post hoc correction. For non-parametric data: Kruskal Wallis test for multiple data sets followed by Dunn’s post hoc test. P values < 0.05 were considered significant. For survival data, Kaplan Meier-survival analysis was used and long-rank Mantel-Cox test was applied to compare two or more survival curves. Pairwise comparisons of individual survival curves were performed manually, and a Bonferroni correction (p value (0.05) / number of comparisons) applied to determine the new threshold of significance.

## Supporting information

Combined supplemental figures

## Data availability statement

All data were stored on the internet server of The Francis Crick Institute and can be requested from the corresponding author.

## Acknowledgments

This project was supported by Cancer Research UK (C422/A29942). M.W. is also supported by the Francis Crick Institute, which receives its core funding from Cancer Research UK (CC2096), the UK Medical Research Council (CC2096), and the Wellcome Trust (CC2096). I.A.M. also acknowledges support from Ovarian Cancer Action (grant number PSN418). For the purpose of open access, the authors have applied a CC BY public copyright licence to any author accepted manuscript version arising from this submission.

## Author contributions

**S.D.:**Conceptualization, data curation, formal analysis, investigation, writing - original draft. **C.J.Q.:** Data curation, formal analysis, investigation, methodology, writing – review & editing. **K.E.T.:** *In vivo* investigation, methodology & validation. **L.A.S.:** Conceptualization, writing – review & editing. **A.P.:** Conceptualization, writing – review & editing. **I.D.R.:** Supervision, formal analysis, methodology, writing – review & editing. **D.P.E.:**Formal analysis **M.H.:**Supervision, resources, methodology. **I.A.M.:**Conceptualization, funding acquisition, resources, supervision, writing – review & editing. **M.W.:** Conceptualization, funding acquisition, resources, supervision, writing – review & editing.

