## Supplementary material for "Vinorelbine enhances the efficacy of GM-CSF-armed oncolytic vaccinia virus in a preclinical model of ovarian high grade serous carcinoma": Combined supplemental figures

A

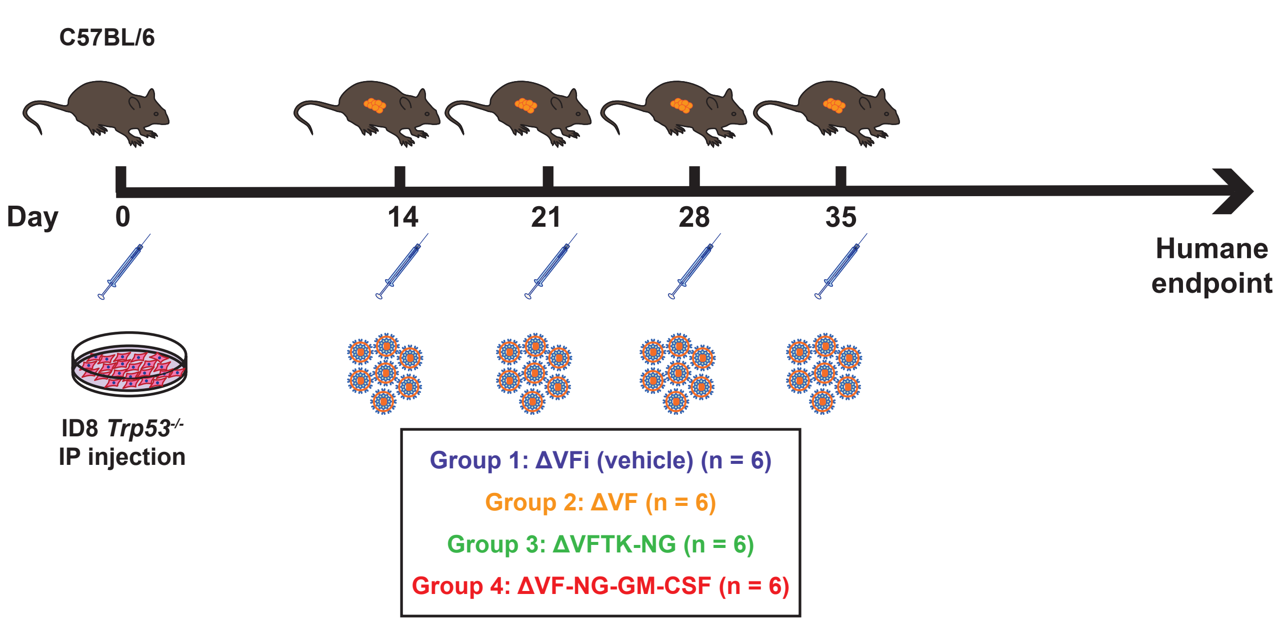


B

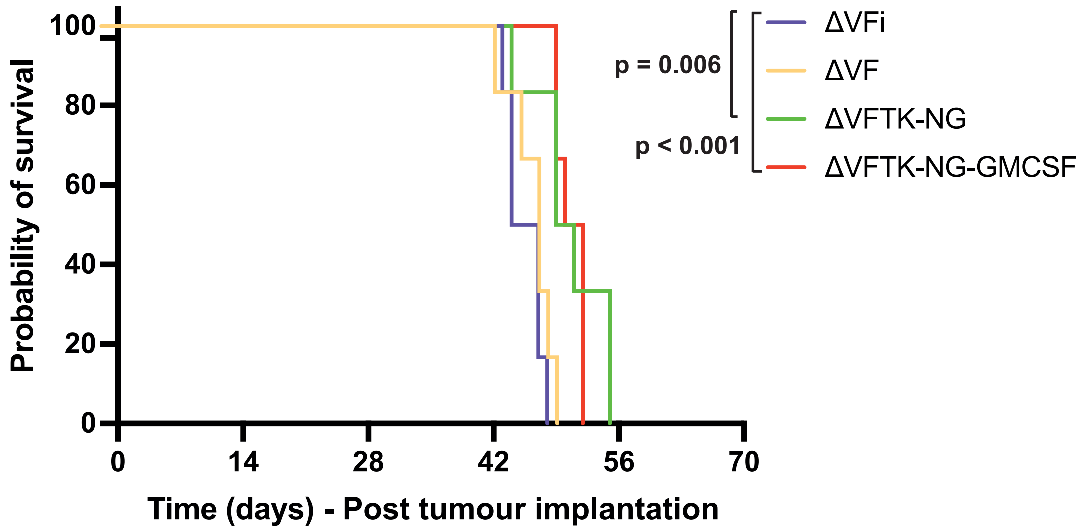


C

| | ΔVFi | ΔVF | ΔVFTK-NG | ΔVFTK-NG-GM-CSF |
|---|---|---|---|---|
| **Omental tumour (mg)** | 112.0 ± 13.0 | 93.0 ± 8.0 | 100.0 ± 3.8 | 89.0 ± 6.6 |
| **Spleen (mg)** | 365.0 ± 31.0 | 274.0 ± 31.0 | 306.0 ± 15.0 | 310.0 ± 40.0 |
| **Liver (mg)** | 848.0 ± 61.0 | 839.0 ± 39.0 | 759.0 ± 45.0 | 808.0 ± 33.0 |
| **Ascites (ml)** | 4.7 ± 0.5 | 3.0 ± 0.5 | 5.4 ± 0.7 | 5.6 ± 0.8 |

**Figure S1. ΔVFTK-NG-GM-CSF provides superior survival benefit in monotherapy** **A**. Schematic representation of the experimental design of the *in vivo* survival study. Four groups of mice were injected IP with ID8 Trp53$^{-/-}$ cells on day 0 and subsequently inoculated with the indicated viruses on day 14, 21, 28 and 35. ΔVFi is the control heat inactivated virus. **B**. Kaplan-Meier survival curve showing survival data for each virus analysed by log-rank test. **C**. Quantification of omental tumour, spleen and liver weights as well as ascitic volumes for each group. Data are represented as mean ± SD. One-way ANOVA was used to determine significance between all groups with Tukey multiple comparisons post-hoc test.

| | 1 | 2 | 3 | 4 | 5 | 6 | 7 | 8 | 9 | 10 | 11 | 12 | 13 | 14 | 15 | 16 | 17 | 18 | 19 | 20 | 21 | 22 | 23 | 24 |
|---|---|---|---|---|---|---|---|---|---|---|---|---|---|---|---|---|---|---|---|---|---|---|---|---|
| A | DMSO | DMSO | AG-1478 | Ellipticine (hydrochloride) | TRAM-34 | Nolatrexed dihydrochloride | Sitagliptin | MMAF-Ome | Evacetrapib | Erastin | Ceftazidime | MMAD | Enzastaurin | Cytochalasin E | TPCA-1 | Aminopterin | IMD-0354 | DMSO | Sepantronium (bromide) | DMSO | Olaparib | DMSO | DMSO | DMSO |
| B | DMSO | DMSO | DMSO | DMSO | DMSO | DMSO | DMSO | DMSO | DMSO | DMSO | DMSO | DMSO | DMSO | DMSO | DMSO | DMSO | DMSO | DMSO | DMSO | DMSO | DMSO | DMSO | DMSO | DMSO |
| C | DMSO | DMSO | Gemcitabine | Geldanamycin | Iniparib | Idasanutlin | Finasteride | SNX-2112 | Chromomycin A3 | MMAF (hydrochloride) | CCT 137690 | DMSO | Methotrexate | DMSO | Alvespimycin (hydrochloride) | Pemetrexed (disodium) | Cabozantinib | DMSO | Pralatrexate | DMSO | Sumatriptan (succinate) | DMSO | DMSO | DMSO |
| D | DMSO | DMSO | DMSO | DMSO | DMSO | DMSO | DMSO | DMSO | DMSO | DMSO | DMSO | DMSO | DMSO | DMSO | DMSO | DMSO | DMSO | DMSO | DMSO | DMSO | DMSO | DMSO | DMSO | DMSO |
| E | DMSO | DMSO | Bortezomib | Harringtonine | Mitoxantrone | AMG131 | GW842166X | Imidazole ketone erastin | PF-573228 | Thapsigargin | NSC319726 | DMSO | Raltitrexed | DMSO | Omipalisib | Dicloxacillin (Sodium hydrate) | CHIR-124 | DMSO | Vinorelbine (ditartrate) | DMSO | Erlotinib | DMSO | DMSO | DMSO |
| F | DMSO | DMSO | DMSO | DMSO | DMSO | DMSO | DMSO | DMSO | DMSO | DMSO | DMSO | DMSO | DMSO | DMSO | DMSO | DMSO | DMSO | DMSO | DMSO | DMSO | DMSO | DMSO | DMSO | DMSO |
| G | DMSO | DMSO | Clotrimazole | Trifluridine | Tolnaftate | Combretastatin A4 | Asimadoline (hydrochloride) | Auristatin F | JNJ-42165279 | PU-H71 | Flucloxacillin sodium | DMSO | Onalespib | DMSO | Vorinostat | DMSO | GSK2334470 | DMSO | TUG-469 | DMSO | CCT241736 | DMSO | DMSO | DMSO |
| H | DMSO | DMSO | DMSO | DMSO | DMSO | DMSO | DMSO | DMSO | DMSO | DMSO | DMSO | DMSO | DMSO | DMSO | DMSO | DMSO | DMSO | DMSO | DMSO | DMSO | DMSO | DMSO | DMSO | DMSO |
| I | DMSO | DMSO | Ruboxistaurin | NVP-2 | Dutasteride | Prexasertib | Daunorubicin (hydrochloride) | Rocaglamide | LY2922470 | WR99210 | Amsacrine (hydrochloride) | Ergosterol | Rosiglitazone | DMSO | AT7519 | DMSO | Canertinib (dihydrochloride) | DMSO | Fasiglifam | DMSO | Nutlin-3a | DMSO | DMSO | DMSO |
| J | DMSO | DMSO | DMSO | DMSO | DMSO | DMSO | DMSO | DMSO | DMSO | DMSO | DMSO | DMSO | DMSO | DMSO | DMSO | DMSO | DMSO | DMSO | DMSO | DMSO | DMSO | DMSO | DMSO | DMSO |
| K | DMSO | DMSO | Tanespimycin | BRL 54443 | Ganetespib | Podofilox | GSK3787 | Abexinostat | SNS-032 | Carfilzomib | Elesclomol | ZSTK474 | MA-0204 | DMSO | BX795 | DMSO | Tubulin inhibitor 1 | DMSO | Torcetrapib | DMSO | JNJ-10198409 | DMSO | DMSO | DMSO |
| L | DMSO | DMSO | DMSO | DMSO | DMSO | DMSO | DMSO | DMSO | DMSO | DMSO | DMSO | DMSO | DMSO | DMSO | DMSO | DMSO | DMSO | DMSO | DMSO | DMSO | DMSO | DMSO | DMSO | DMSO |
| M | DMSO | DMSO | Roflumilast | Diacetoxyscirpenol | Cytarabine | NVP-HSP990 | Trimebutine (maleate) | Silvestrol | Apremilast | Nonactin | CP-91149 | FL118 | Ceftiofur | DMSO | Floxuridine | DMSO | (R)-CR8 | DMSO | NVP-TAE 226 | DMSO | Alogliptin | DMSO | DMSO | DMSO |
| N | DMSO | DMSO | DMSO | DMSO | DMSO | DMSO | DMSO | DMSO | DMSO | DMSO | DMSO | DMSO | DMSO | DMSO | DMSO | DMSO | DMSO | DMSO | DMSO | DMSO | DMSO | DMSO | DMSO | DMSO |
| O | DMSO | DMSO | PF-3845 | CP-316819 | Bay 59-3074 | Diosgenin | Vindesine (sulfate) | Aleglitazar | Seralutinib | SPDB-DM4 | Pazopanib (Hydrochloride) | CH5138303 | 5-Fluorouridine | DMSO | Vinblastine (sulfate) | DMSO | PF-04691502 | DMSO | PS-1145 | DMSO | Dinaciclib | DMSO | DMSO | DMSO |
| P | DMSO | DMSO | DMSO | DMSO | DMSO | DMSO | DMSO | DMSO | DMSO | DMSO | DMSO | DMSO | DMSO | DMSO | DMSO | DMSO | DMSO | DMSO | DMSO | DMSO | DMSO | DMSO | DMSO | DMSO |

Solvant Water
Solvant Ethanol
Solvant DMSO

Red: 2 mM concentration
Black: 10 mM concentration

**Figure S2. Secondary screen mother plate layout**

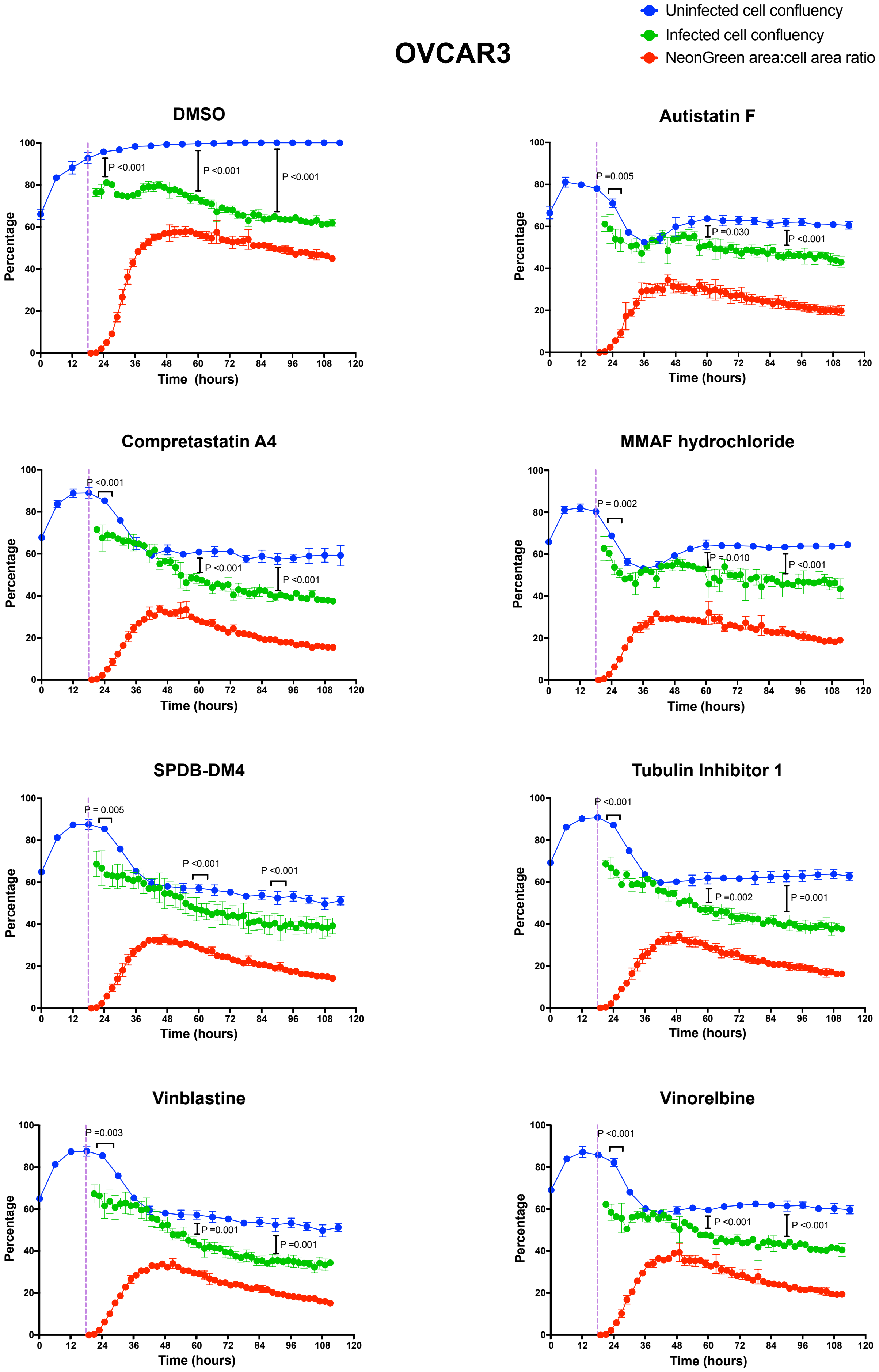


**Figure S3. Tubulin assembly inhibitors enhance vaccinia-induced OVCAR3 cell death** Representative graphs of the combination of the indicated tubulin polymerisation inhibitors with vaccinia in OVCAR3 cell line. The purple dotted line represents infection with ΔVFTK-NG (MOI 0.5) at 18 hours post cell seeding. All compounds were used at 1 μM. Student's t-test was used to compare uninfected against infected cell confluency at 25, 60 and 90 hours post cell seeding. Error bars represent SD.

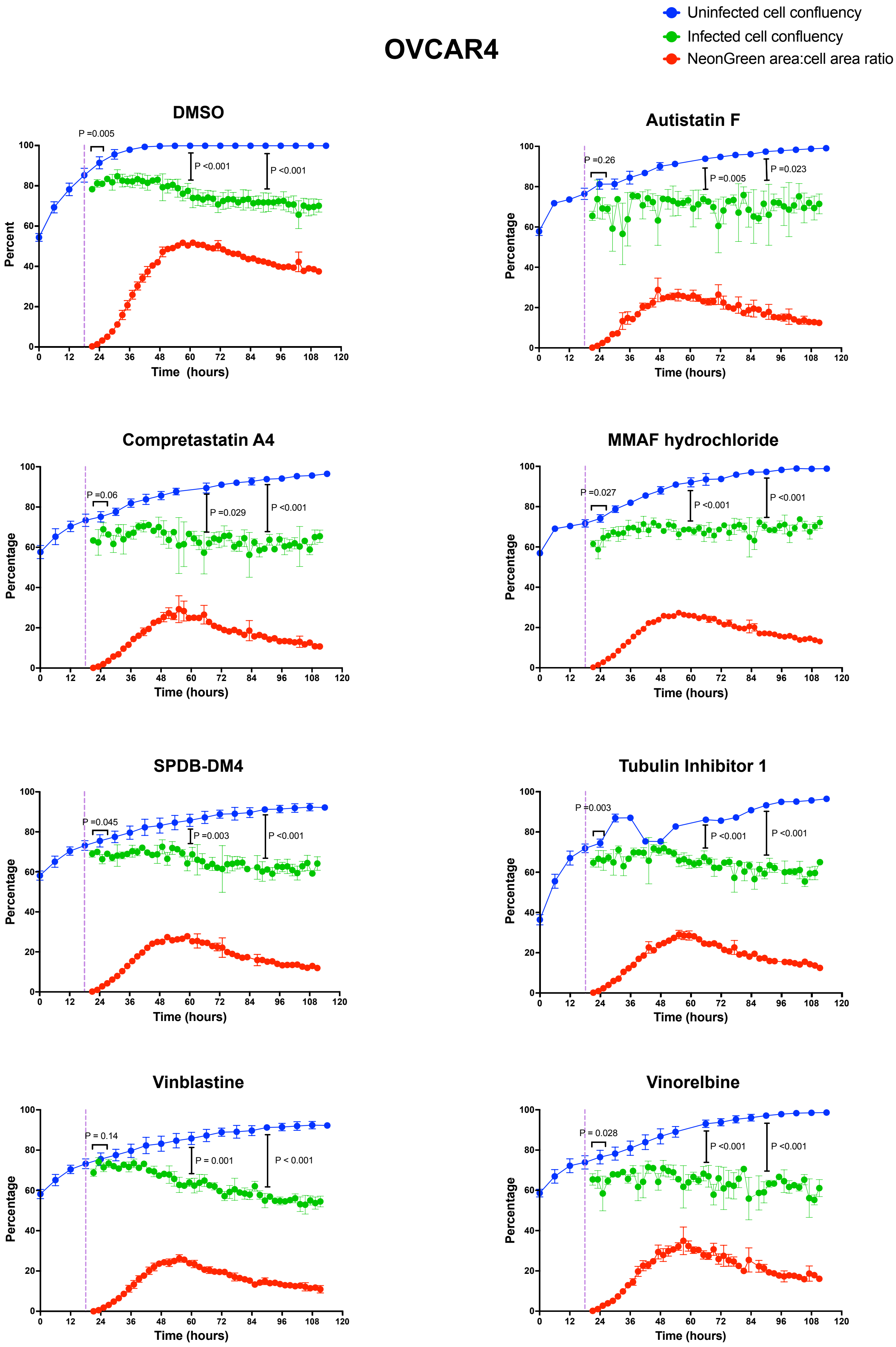


**Figure S4. Tubulin assembly inhibitors enhance vaccinia-induced OVCAR4 cell death** Representative graphs of the combination of the indicated tubulin polymerisation inhibitors with vaccinia in OVCAR cell line. The purple dotted line represents infection with ΔVFTK-NG (MOI 0.5) at 18 hours post cell seeding. All compounds were used at 1 μM. Student's t-test was used to compare uninfected against infected cell confluency at 25, 60 and 90 hours post cell seeding. Error bars represent SD.

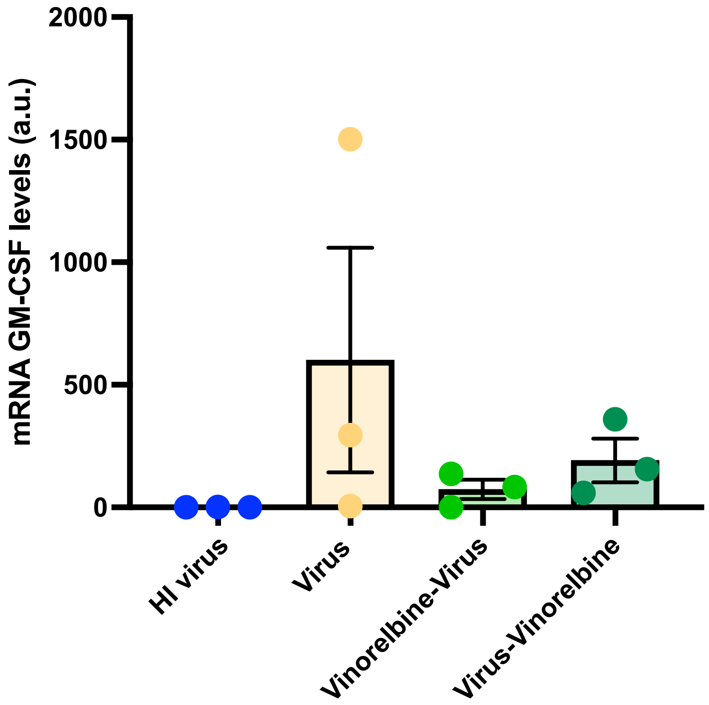


**Figure S5. Quantification of GM-CSF mRNA in ΔVFTK-NG-GM-CSF infected tumours** Quantification of GM-CSF mRNA in omental tumours infected with virus (ΔVFTK-NG-GM-CSF) with or without vinorelbine treatment. HI represents the heat inactivated control virus control. The tumour samples are from the same experiments shown in Figure 5A. Error bars represent mean ± SD.

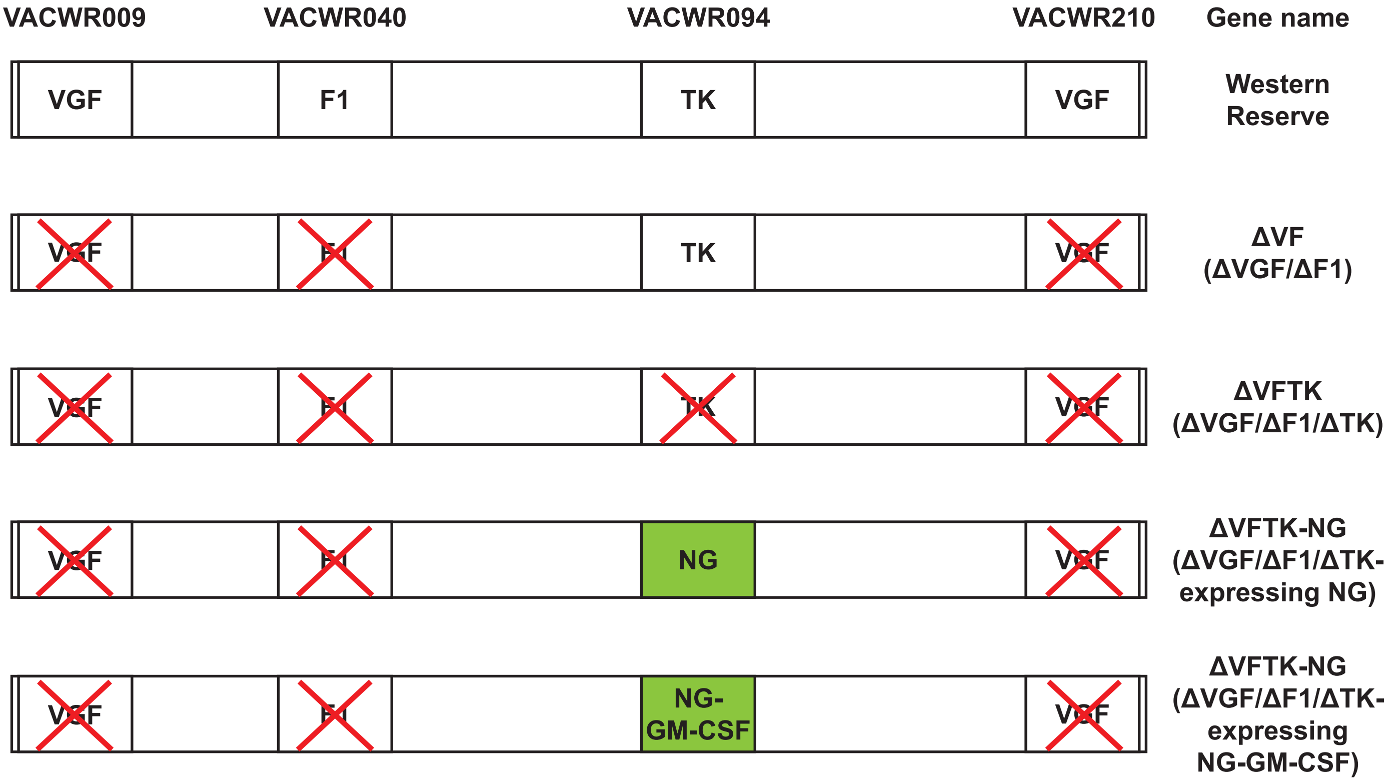


**Figure S6. Generation of recombinant viruses**
Schematic illustrating the gene modifications made for the generation if the indicated recombinant viruses
